# Colorectal cancer susceptibility loci as predictive markers of rectal cancer prognosis after surgery

**DOI:** 10.1101/158238

**Authors:** Yue Hu, Jochen Gaedcke, Georg Emons, Tim Beissbarth, Marian Grade, Peter Jo, Meredith Yeager, Stephen Chanock, Hendrik Wolff, Jordi Camps, B. Michael Ghadimi, Thomas Ried

**Affiliations:** Section of Cancer Genomics, Genetics Branch, National Cancer Institute, Bethesda, MD 20892, USA; Department of General, Visceral and Pediatric Surgery, University Medical Center, 37075 Göttingen, Germany; Department of Medical Statistics, University Medical Center, 37075 Göttingen, Germany; Division of Cancer Epidemiology and Genetics, National Cancer Institute, Rockville, MD 20850, USA; Department of Radiation Oncology, University Medical Center, 37075 Göttingen, Germany

**Author notes:** Correspondence to: Thomas Ried, MD. Center for Cancer Research, National Cancer Institute, Building 50, Room 1408, Bethesda, MD 20892-8010.

**Keywords:** colorectal cancer, rectal cancer, SNP, MYC, prognosis, chemoradiotherapy, radiation resistance, rs6983267, rs12953717, rs4464148, rs4925386

## Abstract

**Background:** Colorectal cancer (CRC) is among the leading causes of cancer death. Rectal cancers account for one third of CRC cases. The role and significance of colorectal cancer risk loci in rectal cancer progression has not been investigated.

**Methods:** We generated and explored a dataset from 230 rectal cancer patients by gene expression microarray analysis of cancer samples and matched controls, and SNP arrays of germline DNA.

**Results:** 8q24 (upstream of *MYC*) and 18q21 (in the intron of *SMAD7)*, two of the loci most strongly linked with colorectal cancer risk, as well as 20q13 (in the intron of *LAMA5)*, are tightly associated with the prognosis of rectal cancer patients. For SNPs on 18q21 (rs12953717 and rs4464148) and 20q13 (rs4925386), alleles that correlate with higher risk for the development of colorectal cancer are associated with shorter disease free survival. However, for rs6983267 on 8q24, the low risk allele is associated with a higher risk for recurrence and metastasis after surgery, and importantly, is strongly correlated with the resistance of colorectal cancer cell lines to chemoradiotherapy. We also found that although *MYC* expression is dramatically increased in cancer, patients with higher levels of *MYC* have a better prognosis. The expression of *SMAD7* is weakly correlated with disease free time. Notably, the presence of the 8q24 and 18q21 SNP alleles is not correlated with a change of expression of *MYC* and *SMAD7.* rs4464148, and probably rs6983267 and rs4925386, are linked with overall survival time of patients.

**Conclusions:** SNPs at three colorectal cancer risk loci detect subpopulations of rectal cancer patients with poor prognosis. rs6983267 probably affects prognosis through interfering with the resistance of cancer cells to chemoradiotherapy.

## Materials and Methods

### Patients

Overall, 230 patients were included in this analysis. All patients were staged as locally advanced (cUICCII/III/IV) and treated according to the CAO/ARO/AIO-94 (Sauer *et al*, 2004) or the CAO/ARO/AIO-04 trial (Rodel *et al*, 2012b). Staging included rigid rectoscopy and endorectal ultrasonography, magnetic resonance imaging (MRI) and/or computed tomography as well as patho-histological diagnosis of an adenocarcinoma. Preoperative staging was performed as clinically assessed T-level (T), lymph node status (N) and distant metastases (M). The results were unified as clinical UICC stage (UICC) (Brierley *et al*, 2009).

Staging and treatment was either performed in Göttingen (n=165) or at the five collaborating German Departments of Surgery or Radioncology (n=65). Preoperative chemoradiotherapy (CRT) was applied with a total irradiation dose of 50.4 Gray (28 × 1.8 Gray) and accompanied by either 5-fluorouracil infusion alone (n=128) or in combination with oxaliplatin (n=102). Six weeks after the completion of preoperative CRT, curative surgery (total mesorectal excision) was performed. Standardized histopathological work-up resulted in histopathological parameters of T-level and lymph node status. If distant metastases developed during preoperative CRT a biopsy for histopathological analysis was taken if possible. Four to six weeks after surgery therapy was completed with either 5-FU alone or in combination with oxaliplatin. Recurrence refers to detection of local recurrence or metastasis. The study was approved by the local ethics committee.

### Tumor Biopsies, RNA/DNA Isolation

During rectoscopy several biopsies of tumor and normal mucosa were taken in addition to the diagnostic sample, as well as EDTA-blood as previously described (Gaedcke *et al*, 2012). For gene expression analyses biopsies were immediately stored in RNAlater and kept at 4°C over night. The following day biopsies were frozen and stored at -20°C. Blood was centrifuged (2200 rpm, 10min at 4°C), plasma and the cellular components (leukocyte phase) were aliquoted and stored at -80°C.

Using TRIZOL (Life Technologies, Rockville, MD, USA) RNA was extracted from RNAlater biopsies as previously described (Grade *et al*, 2011). All samples underwent strict quality assessment. Nucleic acid quantity and purity were determined using the NanoDrop spectrophotometer ND1000 (Thermo Fisher Scientific Inc., Waltham, MA, USA), and by a 2100 Bioanalyzer (Agilent Technologies, Santa Clara, CA, USA). For SNP analyses, after informed consent, 7.5 ml of fresh blood was collected, anti-coagulated by EDTA and centrifuged for 10 min at 2000 rpm, and the plasma carefully removed. Samples were stored at -80 °C until isolation. DNA was isolated using the Blood and Tissue DNA midi kit (Qiagen) according to manufacture’s instruction. Briefly, after adding 100 μp of proteinase K to 1 ml of each sample cells were lysed by adding 1.2 ml of Buffer AL, vortexed and incubated for 10 min at 70°C. DNA was bound to a DNeasy spin column and washed. DNA was eluted in water and stored at 20°C until further use. DNA quantity and quality were assessed using Nanodrop and Bioanalyzer.

### Gene expression Microarray Analysis

Microarrays analysis was performed as previously described (Grade *et al*, 2011). Briefly, 600ng of total RNA was amplified and transcribed into fluorescently labeled cRNA following the “Low RNA Input linear Amplification Kit Plus, One Color” protocol (Agilent Technologies, Inc. 2007; Cat. N°C: 5188-5339). cRNA was hybridized to the Human 4 × 44 K v2 array platform from Agilent Technologies (G4845A) as recommended by the manufacturer. Cy3 intensities were detected by one-color scanning using an Agilent DNA microarray scanner (G2505B) at 5 micron resolution. Scanned image files were visually inspected for artifacts and then analyzed using the Agilent feature extraction software (Agilent Technologies, Santa Clara, CA, USA).

The signal intensities from the gene expression arrays were log2 transformed before being normalized to the 75 percentile as instructed by the Agilent protocol. Probes with maximum intensity over all samples of at least 100 were used for further analysis. Gene fold change of each cancer sample was calculated with paired control as reference if available, otherwise the geometric average intensity of all control samples was used as reference. Gene expression data were deposited to Gene Expression Omnibus (GSE87211).

### Illumina Infinium HD assay

High-throughput, genome-wide SNP genotyping, using Infinium Human1M-Quadv1 technology (Part 15003560; Illumina Inc. San Diego, CA), was performed at the Cancer Genomics Research Laboratory (CGR), NCI. Genotyping was performed according to the manufacturer’s guidelines using the Infinium HD Assay automated protocol. Samples were denatured and neutralized then isothermally amplified by whole-genome amplification. The amplified product was enzymatically fragmented, then precipitated and re-suspended before hybridization to the BeadChip. Single-base extension of the oligos on the BeadChip, using the captured DNA as a template, incorporates tagged nucleotides on the BeadChip, which are subsequently fluorophore labeled during staining. The fluorescent label determines the genotype call for the sample. The Illumina iScan scanned the BeadChips at two wavelengths to create image files. Genotypes were called using Illumina GenomeStudio software and exported for analysis.

### Statistical analysis

A change of gene expression with a threshold of 3-fold change (either up or down) and a P value of 0.05 in two tailed unpaired t-test after multiple test correction was considered to be significant. The list of genes, the expression of which significantly differed between the cancer and the mucosa control, was used for Ingenuity Pathway Analysis (IPA) (Qiagen, Hilden, Germany).

PCA (Principal Component Analysis) was conducted utilizing Partek Genomics Suite (Partek Inc, Chesterfield, MO) for log2 transformed intensity (Figure 1A) or log2 fold change (Figure 1B-I). Data points of patient 93 and patient 195 samples were removed in Figure 1B-I to get an extensive view of the whole group.

**Figure 1.**
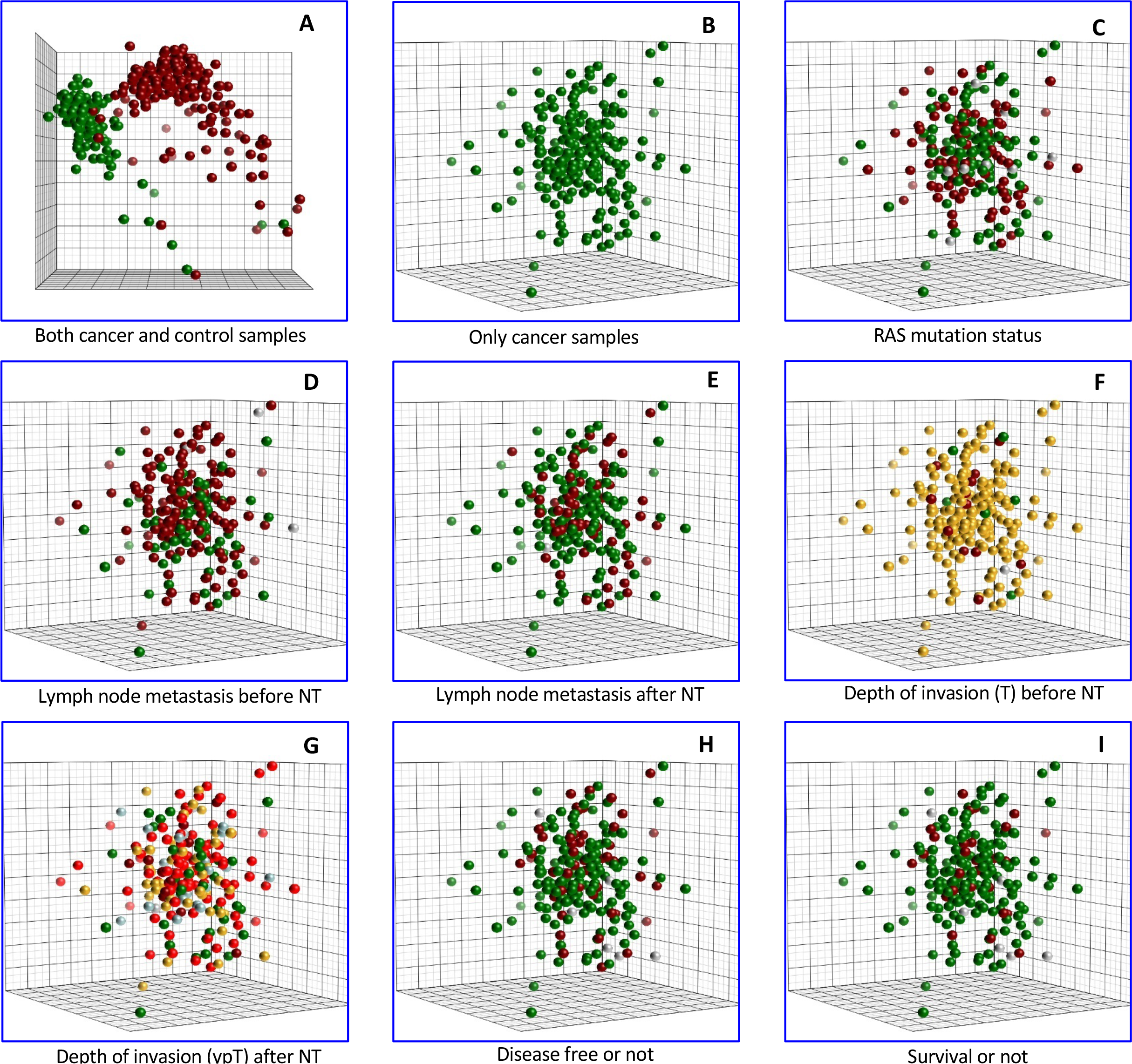
Principal component analysis (PCA) of expression levels of rectal cancer samples and matched normal mucosa. PCA of gene expression of both control samples (green) and rectal cancer samples (red) (A), of expression fold change of rectal cancer samples compared with paired controls (B-I). Samples with (C) RAS mutation (red) or without mutation (green), with lymph node metastasis (red) or without (green) before (D) or after (E) CRT, (F) with depth of invasion before CRT including T2 (green), T3 (yellow) or T4 (crimson), (G) with depth of invasion after CRT including T0 (green), T1 (blue), T2 (yellow), T3 (red) or T4 (crimson), (H) with recurrence after surgery (red) or disease free (green), (I) death due to CRC (red) or survival (green).

Survival curves were constructed using the Kaplan-Meier method and analyzed by the log-rank test. Survival data were evaluated using the univariate and multivariate Cox proportional hazards model. Variables with a significant value of *P* < 0.05 in univariate analysis were used in construction of subsequent multivariate Cox models. Cox *P* value is used unless otherwise stated. Samples were separated into two groups of equal size for high and low gene expression.

The SNPs tested in this study influence CRC occurrence at very different strengths (Broderick *et al*, 2007; Houlston *et al*, 2008; Tomlinson *et al*, 2007). From biological intuition, a factor’s ability to increase CRC risk is likely to be correlated to its potential effect on recurrence risk, and such an inequality will invalidate the assumption of multiple test correction. This inequality can be revealed by enrichment analysis. Among 3 loci with significant effect on rectal cancer recurrence, two are localized on the no.1 and no.3 locus with highest CRC risk in the total list of 21 loci. The biased distribution *P* < 0.028 confirmed the significant association between CRC risk and recurrence risk. With this confirmed inequality, multiple test correction was not applied for SNP number in this study.

## Introduction

Colorectal cancer (CRC) is the third most common cancer and the second leading cause of cancer death in the USA. Each year there are more than 130,000 new cases and more than 49,000 deaths from CRC (Ferlay *et al*, 2015). Rectal cancers account for some 35% of all CRC cases (American Cancer Society, 2016). Despite similarities in the genetic aberrations profiles rectal cancer and colon cancer show significant differences in clinical outcome. Anatomic differences also influence the treatment of these cancers. While colon cancers are usually surgically removed and then treated with adjuvant chemotherapy, the standard treatment for locally advanced rectal cancers consists of a preoperative chemoradiotherapy (CRT) (Rodel *et al*, 2012a; Sauer *et al*, 2004) followed by a total mesorectal excision of the tumor. Furthermore, the typical site of metastasis or progression is different: colon cancers metastasize primarily to liver and have a tendency to spread to the peritoneum, while distal rectal cancers often surpass the liver and spread to the lung (Riihimaki *et al*, 2016). Molecular and genetic studies as well reveal both remarkable similarities but also some noteworthy distinctions. For example, *APC* mutations occur with higher frequency in rectal cancer than in colon cancer (Hong *et al*, 2012; van der Sijp *et al*, 2016).

Many genome-wide association studies (GWAS) have successfully identified more than 20 SNPs associated with the risk for the development of CRC (reviewed in Peters *et al*, 2013; Zhang *et al*, 2014). As outlined above, colon and rectal cancers are very different in their clinical behavior. To our knowledge, the prognostic value of these SNPs has not been systematically evaluated in rectal cancer patients. Here we analyze an extensive dataset of rectal cancers to identify if any of these SNPs are also associated with outcome in rectal cancer patients. SNPs associated with prognosis might augment already established clinical tests and help to individualize a patients’ treatment.

The most extensively studied risk locus of cancer is the SNP rs6983267 on chromosome band 8q24. The G allele of rs6983267 increases the risk of developing several cancers, including CRC, prostate cancer and breast cancer (Haiman *et al*, 2007; Jones *et al*, 2012). Several hypotheses have been proposed to explain the effect of rs6983267, including enhanced binding of the WNT pathway transcription factor TCF7L2 in the *MYC* promoter region, hence activating transcription (Pomerantz *et al*, 2009; Wright *et al*, 2010). Another possible mechanism of action might be interference with the expression of the noncoding RNA CCAT2 and CARLo-5 (Kim *et al*, 2014; Ling *et al*, 2013). The second most studied risk locus on 18q21 contains three SNPs: rs4939827, rs12953717 and rs4464148. They map to introns of the tumor suppressor gene *SMAD7* (Curtin *et al*, 2009; Tenesa *et al*, 2008), influencing its expression. *SMAD7* is involved in the regulation of the TGF-β pathway (Luo *et al*, 2014).

In this study, we explore whether the SNPs that affect the risk of the development of CRC also alter the prognosis of rectal cancer patients and whether their presence affects the expression of adjacent genes.

## Results

### Global gene expression changes are neither associated with survival, lymph node metastasis and depth of invasion, nor with *KRAS* mutation status

The Principal Component Analysis (PCA) of expression data distinguishes cancer samples clearly from control samples, illustrating the drastic change of gene expression patterns during tumorigenesis. As expected, the normal controls cluster more tightly together than the tumor samples (Figure 1A). The list of genes significantly up-regulated in tumors relative to mucosa is enriched for genes associated with WNT-β catenin signaling. Genes significantly down-regulated are linked particularly with various metabolic pathways (Supplementary Table 2). The general expression pattern of cancer samples forms no apparent subgroups (Figure 1B), and there is no separation according to any prognostic parameters, including lymph node metastasis or depth of invasion before or after preoperative CRT, *KRAS* mutation status, disease free survival (DSF) or overall survival (OS) of the patients (Figure 1C-I). Consistently, differential expression analysis for any of those prognostic parameters, e.g., comparison between cancer samples with or without *KRAS* mutations, revealed no statistically significant differences in gene expression (data not shown). Lymph node metastasis and depth of invasion scores from biopsies after, but not before CRT, show a significant association with DSF and OS (Table 2, Supplementary Figure 1A-D, F-I). *KRAS* mutation status serves as an established biomarker for treatment with EGFR inhibitors, such as panitumumab and cetuximab. Only patients with tumors that do not harbor *KRAS* mutations benefit from anti-EGFR therapy (Cunningham *et al*, 2004). Our data show that the *KRAS* mutation status cannot predict response to neoadjuvant chemoradiotherapy (Table 2, Supplementary Figure 1 E, J).

**Table 1.** CRC occurrence risk loci and their association with disease free time and survival time of rectal cancer patients. Loci are ordered according to P value associated with CRC occurrence from Houlston *et al*,(2010) Houlston *et al*, (2008) and Whiffin *et al*, (2014). HR, CI 95% and P value are calculated with univariate Cox proportional hazard model. Significant P values (<0.05) are highlighted with bold. P value and HR calculation for dominant and recessive effect SNPs: rs6983267 TT+GT versus GG, rs4939827 TT+CT versus CC, rs12953717 TT+CT versus CC, rs4925386 TT+CT versus CC; for additive effect SNP rs4925386: TT=0, CT=1, CC=2; for other SNPs: CRC occurrence risky homozygous allele versus non-risky homozygous allele. SNP rs16892766 does not have enough CC allele samples for calculation. SNP rs647161 is not on the SNP array and its data is not available. SNP rs6687758 does not have a death case for the AA reference allele for survival time analysis. See supplementary table 4 for a complete list of P value and HR of CRC disease free time.

| Cytoband | Gene | SNP | CRC occurrence |  | rectal cancer recurrence |  |  |  | rectal cancer survival |  |
| --- | --- | --- | --- | --- | --- | --- | --- | --- | --- | --- |
|  |  |  | Risky allele | No-Risky allele | Risky allele | Genetic effect | HR (95% CI) | *P* | HR (95% CI) | *P* |
| 8q24 | MYC | rs6983267 | G | T | T | dominant | 2.21(1.03-4.74) | **0.042** | 2.61(0.9-7.56) | 0.076 |
| 11q23 | POU2AF1 | rs3802842 | C | A |  |  | 0.94(2.45-0.36) | 0.905 | 0.8(0.26-2.43) | 0.694 |
| 18q21 | SMAD7 | rs12953717 | T | C | T | dominant | 2.46(1.1-5.5) | **0.029** | 1.38(0.56-3.43) | 0.485 |
|  |  | rs4464148 | C | T | C | additive | 1.8(1.19-2.72) | **0.005** | 1.87(1.11-3.13) | **0.019** |
|  |  | rs4939827 | T | C | T | dominant | 2.11(0.83-5.35) | 0.115 | 1.26(0.44-3.66) | 0.668 |
| 8q23 | EIF3H | rs16892766 | A | C |  |  | --- | --- | --- | --- |
| 15q13 | GREM1 | rs4779584 | T | C |  |  | 1.78(0.54-5.87) | 0.346 | 3(0.87-10.42) | 0.083 |
| 19q13 | RHPN2 | rs10411210 | C | T |  |  | 1.35(0.19-9.84) | 0.767 | 0.84(0.11-6.25) | 0.864 |
| 20p12 | BMP2 | rs961253 | A | C |  |  | 0.8(1.86-0.34) | 0.6 | 0.56(1.92-0.16) | 0.356 |
| 16q22 | CDH1 | rs9929218 | G | A |  |  | 0.46(1.13-0.19) | 0.089 | 0.95(4.22-0.22) | 0.951 |
| 14q22 | BMP4 | rs4444235 | C | T |  |  | 1.12(0.47-2.64) | 0.796 | 0.88(0.25-3.14) | 0.849 |
| 11q13 | POLD3 | rs3824999 | G | T |  |  | 1.19(0.55-2.6) | 0.658 | 0.69(0.23-2.07) | 0.512 |
| 10p14 | FLJ3802842 | rs10795668 | G | A |  |  | 2.44(0.58-10.37) | 0.226 | 3.52(0.45-27.42) | 0.229 |
| 20q13 | LAMA5 | rs4925386 | C | T | C | recessive | 0.43(0.24-0.79) | **0.007** | 0.46(0.21-1.01) | 0.053 |
| 20p12 | BMP2 | rs4813802 | G | T |  |  | 1.18(0.46-2.99) | 0.729 | 1.38(0.48-3.96) | 0.555 |
| 12q12 | DIP2 | rs11169552 | C | T |  |  | 1.97(0.47-8.3) | 0.357 | 1.27(0.29-5.59) | 0.755 |
| 12q12 | ATF1 | rs7136702 | T | C |  |  | 1.1(0.4-3.03) | 0.85 | 0.31(0.04-2.46) | 0.27 |
| 3q36 | TERC | rs10936599 | C | T |  |  | 1.65(0.39-6.97) | 0.498 | 2.29(0.3-17.53) | 0.426 |
| 20p12 | HAO1 | rs2423279 | C | T |  |  | 0.8(2.31-0.28) | 0.679 | 0.99(3.4-0.29) | 0.983 |
| 1q41 | DUSP10 | rs6691170 | T | G |  |  | 1.16(0.5-2.69) | 0.723 | 0.84(0.32-2.2) | 0.718 |
| 1q41 | DUSP10 | rs6687758 | G | A |  |  | 0.85(3.6-0.2) | 0.827 | --- | --- |
| 6p21 | CDKN1A | rs1321311 | A | C |  |  | 0.5(3.66-0.07) | 0.491 | 1.66(12.95-0.21) | 0.631 |
| 5q31 | PITX1 | rs647161 | A | C |  |  | --- | --- | --- | --- |
| 12p13 | CCND2 | rs10774214 | T | C |  |  | 1.33(0.64-2.78) | 0.447 | 0.96(0.36-2.52) | 0.929 |

**Table 2.**
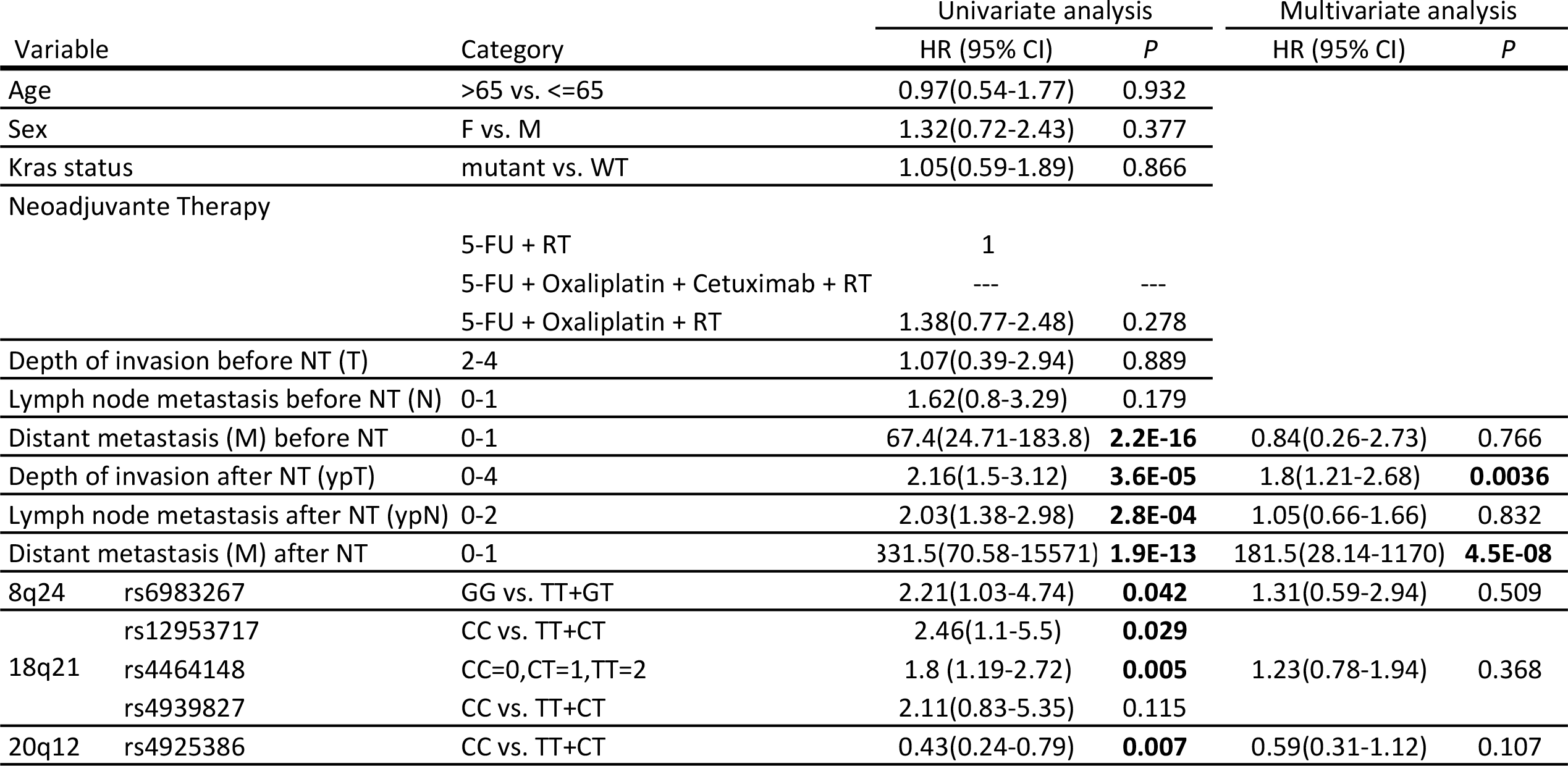
Univariate and multivariate analyses of SNPs and clinicophathological variables for disease free time (Cox proportional hazards regression) Because of high linkage disequilibrium among 18q21 SNPs, only rs4464148 is included in multivariate analysis. Significant P values (<0.05) are highlighted in bold.

### Association of polymorphisms and interval to recurrence after surgery

Many GWAS of CRC have identified polymorphisms in at least 22 loci associated with an increased risk of developing CRC. We investigated the association of 21 of those loci with DSF and OS. We found three loci that are significantly associated with rectal cancer recurrence, including rs6983267 on 8q24 upstream of *MYC*, rs12953717 and rs4464148 on 18q21 in an intron of *SMAD7*, and rs4925386 on 20q13 in an intron of *LAMA5* (Table 1). The SNP rs4464148 on 18q21 also has a significant effect on OS, while the SNP rs6983267 on 8q24 (P = 0.076) and the SNP rs4925386 on 20q13 (P = 0.053) have noticeable effect although not significant (Table 1). 8q24 and 18q21 are among the loci with the most significant P value for CRC risk (Table 1) (Whiffin *et al*, 2014). Interestingly, rs6983267 has opposite effects when analyzed for CRC susceptibility and prognosis. The high-risk allele G of rs6983267 correlates with an increase in CRC incidence in the population, but is associated with a longer DSF and OS for rectal cancer patients when compared to the low-risk allele T (Table 1). We have investigated the difference in sensitivity of CRC cell lines to CRT (Spitzner *et al*, 2014). We found that the CRT sensitivities of CRC cell lines are tightly correlated with their rs6983267 genotypes. Cell lines with higher CRT resistance invariably carry the T allele that is associated with short DSF in rectal cancer patients (Figure 2F). Such an allelic specific distribution is not observed for any other significant SNPs (Supplementary Figure 2).

**Figure 2.**
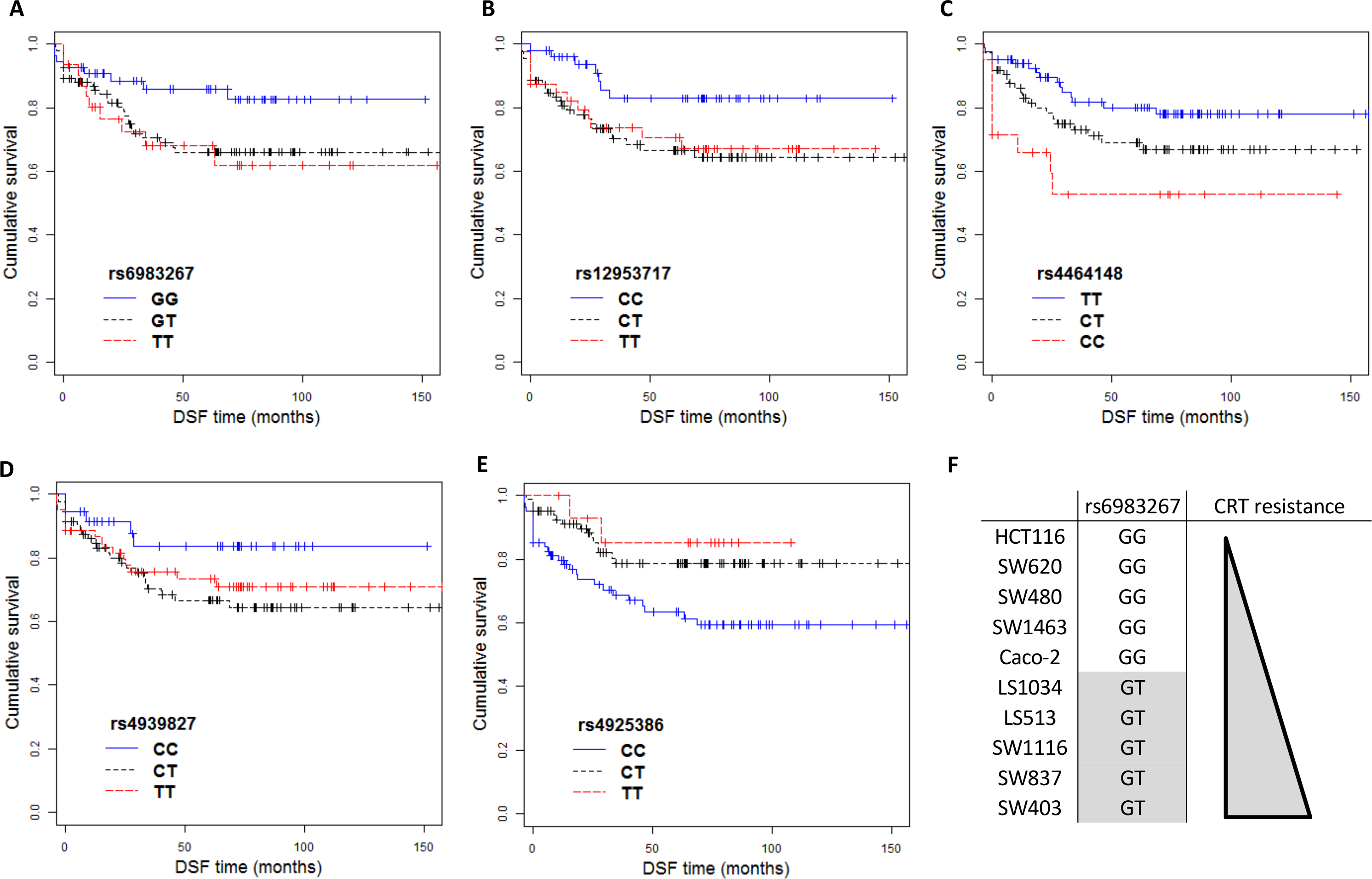
Influence of CRC development associated SNPs on DFS of rectal cancer patients and on CRT resistance of CRC cell lines. Kaplan-Meier curves of DSF of 8q24 SNP rs6983267 (A), 18q21 SNPs rs12953717 (B) rs4464148 (C) and rs4939827 (D), 20q13 SNP rs4925386 (E) for rectal cancer patients. (F) Genotype of rs6983267 for CRC cell lines. Cell lines are ordered according to their CRT resistance as in Spitzner *et al*, (2014) Figure C. Genotypes unfavorable to rectal cancer patients are highlighted with gray. See supplementary figure 2 for data of all 5 SNPs.

Allele G of rs6983267 enhances CRC incidence with a dosage effect; this fits with an additive model (Li *et al*, 2008), while CRC recurrence of patients carrying the allele T fits best with a dominant model; thus, patients with the GT allele have survival curves very similar to the TT allele (DSF: HR = 2.21, 95% CI 1.03-4.74, *P* = 0.042) (Figure 2A). In contrast, the high-risk alleles for CRC occurrence and recurrence are the same for rs12953717 and rs4464148 (Table 1). The 18q21 SNP rs12953717 (DSF: HR = 2.46, 95% CI 1.11-5.5; *P* = 0.039) and the 20q13 SNP rs4925386 (DSF: HR = 0.43, 95% CI 0.24-0.79; *P* = 0.007) fit best with the dominant and recessive model, respectively (Figure 2B,E), while the 18q21 SNP rs4464148 fits best with the additive model for both DSF and OS, with a 1.8-fold hazard risk for DSF (95% CI 1.19-2.72; *P* = 0.005) (Figure 2C) and 1.87-fold hazard risk for OS (95% CI 1.11-3.13; *P* = 0.019) (Supplementary Figure 4C) for each high-risk allele. Multivariate Cox analysis shows only depth of invasion (ypT) and distant metastasis (M) after CRT as significant predictors of DSF (Table 2).

**Figure 3.**
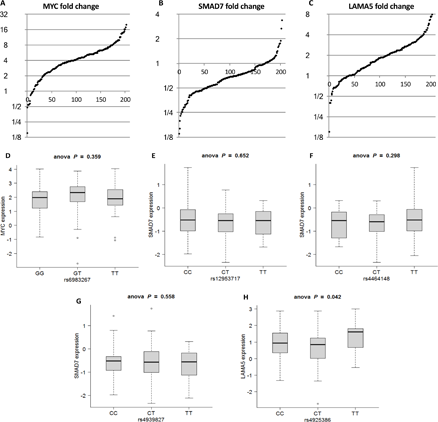
Expressions levels of *MYC, SMAD7* and *LAMA5* have no association with adjacent significant SNPs. Fold change of expression of *MYC* (A), *SMAD7* (B) and *LAMA4* (C) of rectal cancer samples comparing with paired controls. Expression of *MYC* in cancer samples with different alleles of rs6983267 (D), *SMAD7* with different alleles of rs12953717 (E) rs4464148 (F) and rs4939827 (G) and *LAMA5* with different alleles of rs4925386 (H). One way Anova P value was calculated.

**Figure 4.**
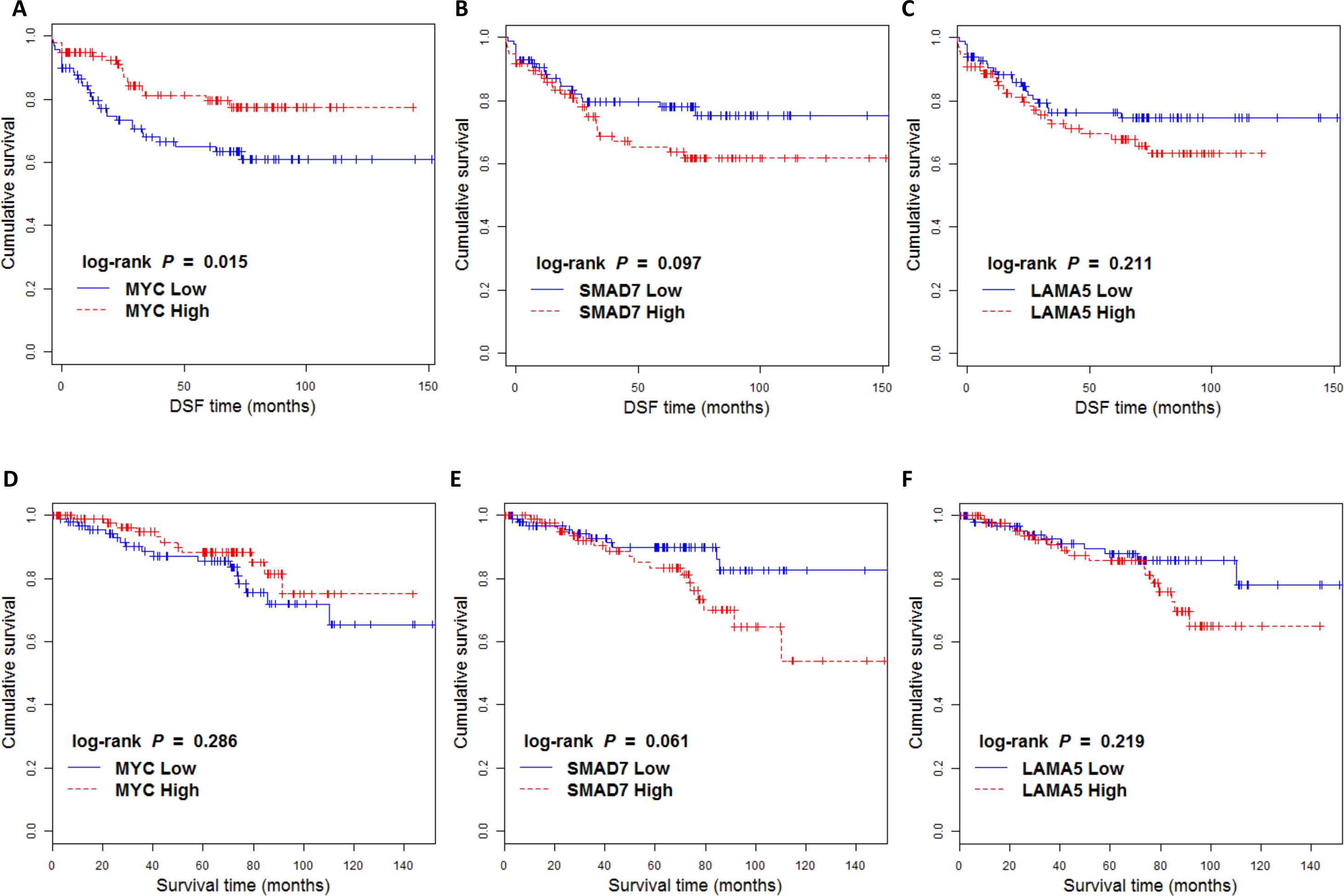
Low *MYC* expression is significantly associated with disease free time of rectal cancer patients. Kaplan-Meier curves of disease free time of high and low expression of *MYC* (A), *SMAD7* (B) and *LAMA5* (C). Kaplan-Meier curves of survival of high and low expression of *MYC* (D), *SMAD7* (E) and *LAMA5* (F).

After multiple test correction, we did not detect any significantly differentially expressed genes among samples with different rs6983267 genotypes (data not shown). However, we discovered that, under a loose criterion with no correction for multiple testing, the expression of two radioresistance related genes, *NOS1* and *AATK*, is correlated with allele number and has a P value < 0.05 between homozygous alleles (Supplementary Figure 3). Cells derived from *NOS1* homozygous knockout mouse have significantly enhanced radioresistance (Epperly *et al*, 2007; Rajagopalan *et al*, 2010). *AATK* siRNA knockdown promotes cell line survival after ionizing radiation while its overexpression reverses this effect (Zhu *et al*, 2016). In line with these reports, the expression of both *NOS1* and *AATK* have the lowest expression in samples with TT alleles, intermediate expression in samples with GT alleles, and highest expression in samples with GG alleles.

### Association between prognosis and the expression of genes adjacent to SNPs associated with CRC

We examined the expression of genes potentially linked with the SNPs on 8q24, 18q21 and 20q13, which are MYC, *SMAD7* and *LAMA5*, respectively. Our data show no correlation between their expression levels and the presence of specific SNPs (Figure 3D-H). In addition, the overall expression levels of these genes with respect to different alleles for any of these SNPs are indistinguishable from each other (data not shown). But *MYC*, and to a lesser extent *SMAD7*, do have an association with prognosis. Compared to matched normal mucosa, rectal cancers have significantly higher expression of *MYC* and lower expression of *SMAD7* (Figure 3A-B). However, patients with lower *MYC* expression or higher *SMAD7* expression have shorter DSF and probably shorter OS, though the P value is only significant for effect of *MYC* expression on DSF (HR = 0.49, 96% CI 0.27-0.88; *P* = 0.017) (Figure 4A-B). *LAMA5* expression is consistently increased in cancer samples (Figure 3C), but it is not correlated with prognosis (Figure 4C, 4F). These results were confirmed with independent CRC sample sets (Supplementary Figure 5).

## Discussion

Previous GWAS studies have identified dozens of loci associated with an increased risk for the development of CRC (Peters *et al*, 2013; Zhang *et al*, 2014). In this study we determined whether those loci affect the prognosis of patients with rectal cancers, which comprise approximately 35% of patients with CRC (Society, 2016). We did observe that three loci, on chromosome bands 8q24, 18q21 and 20q13, significantly influence DSF, and to a lesser extent, OS. It is reasonable to assume that genetic factors that increase the risk for cancer development could also affect disease prognostication. Consistently, we found that two out of these three loci (8q24 and 18q21) are among the loci with strongest association with CRC risk (significance rank no.1 and no.3 respectively in the GWAS study of Houlston 2008). They are also the 1^st^ and 2^nd^ loci found associated with colorectal cancer (Broderick *et al*, 2007; Houlston *et al*, 2008; Tomlinson *et al*, 2007). Both loci are associated with highly significant P value in many GWAS (Peters *et al*, 2015) and are the two most extensively studies studied CRC risk loci.

rs6983267 on 8q24 has been found to be strongly associated with multiple cancer types, including CRC (Haiman *et al*, 2007), prostate cancer (Yeager *et al*, 2007), and thyroid cancer (Jones *et al*, 2012). Interestingly, 8q24 SNP rs6983267 has divergent effects on CRC occurrence and prognosis. This is reminiscent of the effect of *BRAC1/2* mutations on breast cancer risk, which is significantly increased in women carrying these mutations. But they correlate with longer DSF and OS time in patients with triple negative breast cancers (Anders *et al*, 2013). Such a reverse association of rs6983267 is also found in prostate cancer, where the G allele correlates with the risk of cancer occurrence, while the T allele is significantly associated with metastatic prostate cancer, with 1.49 risk ratio over the G allele (Ahn *et al*, 2011). The fact that rectal cancer and prostate cancer share such a phenomenon suggests a common mechanism. One possible mechanism is that the G allele of rs6983267 lowers the threshold for genetic changes that result in CRC occurrence, such that patients with the G allele tend to accumulate fewer mutations and genomic imbalances when the tumor is detectable, and thus have a better prognosis. It will be interesting to see if this phenomenon holds true in the other cancer types influenced by rs6983267, such as thyroid and breast cancers.

Intriguingly, the CRT resistance levels of CRC cell lines are tightly correlated with the genotype of rs6983267 SNP such that cell lines with homozygous GG alleles are always more CRT sensitive than cell lines carrying GT alleles. This discovery suggests that the apparent adverse effect of the T allele in rectal and prostate cancer patients might be due to its role in increasing the CRT resistance of cancer cells. Radiosensitivity related genes with allelic differential expression such as *NOS1* and *AATK* might contribute to this effect (Supplementary Figure 3). This effect might also arise from allele-related differences in general expression response to CRT, which could be revealed by a future meta-analysis of post-CRT samples.

Rectal cancer samples have greatly increased expression levels of *MYC* compared to matched normal mucosa, while the expression of *SMAD7* is consistently decreased; however, patients with higher *MYC* expression or lower *SMAD7* expression show a better prognosis than patients with an opposite expression pattern (see below).

The SNPs on 18q21 locus also correlates with prognosis. But unlike rs6983267, the alleles on 18q21, rs4939827, rs12953717 and rs4464148 have the same effect on CRC risk and recurrence of patients with rectal cancer. These three SNPs are in the same intron of *SMAD7* and are in high linkage disequilibrium (Li *et al*, 2008; Thompson *et al*, 2009). We found that the alleles of rs12953717 and rs4464148 are significantly associated with DSF. The 3^rd^ SNP, rs4939827, shows the same trend, although with an insignificant P value, likely due to the limited sample size (Figure 2B, Table 1). Not surprisingly, the directions of the effects on DSF are the same as on OS for those SNPs affecting both DSF and OS (rs4464148, rs6983267 and rs4925386).

The three loci correlated with prognosis of rectal cancer patients are adjacent to *MYC, SMAD7* and *LAMA5*, respectively. *MYC* is involved both in hematological malignancies and solid tumors, and is required for oncogenesis in a mouse model of CRC (Ignatenko *et al*, 2006; Sansom *et al*, 2007). *MYC* is also an important regulator of cancer stem cells (Curry *et al*, 2015). Cancer stem cells are significantly more resistant to CRT than the bulk tumor and are thought to be responsible for cancer recurrence after therapy (Krause *et al*, 2017). One conceivable mechanism is that the long-term effect of rs6983267 on *MYC* expression perturbs the stemness of cells and results in a differential response of cancer cells to CRT. We found that the majority of CRC cancer samples have dramatically higher expression of *MYC* (>80% samples with >2 fold increase and >50% with >4 fold increase Figure 3F), probably due to aberrant Wnt/β-catenin signaling (Rennoll & Yochum, 2015). Surprisingly, again we found that the expression level of *MYC* is reversely correlated with tumor aggressiveness; patients with high *MYC* expression having a longer DSF after surgery. rs6983267 has been reported to enhance binding of the transcription factor TCF7L2 to the enhancer of *MYC* (Rosse *et al*, 2014; Verzi *et al*, 2010). One could hypothesize that rs6983267 exerts its effect on CRC by directly interfering with *MYC* expression level. However, the expression level of *MYC* in both mucosa and tumor tissues is not correlated with rs6983267 alleles (Figure 3A), even when normalized to genomic copy numbers of this locus (data not shown). This result does not rule out the possibility that the presence of rs6983267 influences CRC risk via an alteration of *MYC* expression during a transient but important period of tumorigenesis, but this change, if it exists, is masked by other process at later stages of CRC that drastically increase *MYC* expression.

*SMAD7* is involved in tumorigenesis and metastasis (Halder *et al*, 2005; Halder *et al*, 2008). The samples analyzed in the present data set show dramatically decreased levels of *SMAD7* expression when compared to matched normal mucosa. *SMAD7* expression levels showed a weak correlation with prognosis (Figure 3J, Supplementary Figure 5B). Intriguingly, patients with lower *SMAD7* expression have a longer DSF. The result is consistent with the finding that loss of *SMAD7* is associated with a favorable outcome for CRC patients, while amplification of *SMAD7* worsens survival (Boulay *et al*, 2003). Therefore, for both *MYC* and *SMAD7*, the change in expression in cancer comparing with control tissue is reversely correlated with DSF. Our data show that there is no direct correlation between the alleles of SNPs and the expression of these genes in tumor samples (Figure 3A-D) or control samples (data not shown). In addition, the presence of different alleles (high-risk or low-risk) does not determine global gene expression levels (data not shown). It therefore remains to be determined on how precisely these SNPs influence cancer risk.

In summary, our results suggest that polymorphisms at specific SNPs on chromosome bands 8q24, 18q21 and 20q13 hold promise as clinically useful tools to predict therapy outcomes and long term follow-up of rectal cancer patients. It will be informative to study the correlation between these SNPs and other CRC prognostic factors and to study whether there exists a differential response to various treatments used in patients with CRC depending on the presence of distinct alleles. These studies will provide a deeper understanding of the molecular mechanisms underlying the idiosyncrasies of individual tumor behavior and disease outcome, and hopefully will help to develop evidence-based individualized treatments.

## Acknowledgements

This study was supported in part by the Intramural Research Program, NCI/NIH. The authors are grateful to Dr. Reinhard Ebner for critical comments on the manuscript.

## References

AhnJ, KibelAS, ParkJY, RebbeckTR, RennertH, StanfordJL, OstranderEA, ChanockS, WangMH, MittalRD, IsaacsWB, PlatzEA, HayesRB (2011) Prostate cancer predisposition loci and risk of metastatic disease and prostate cancer recurrence. Clin Cancer Res 17(5): 1075–1081

AndersCK, ZagarTM, CareyLA (2013) The management of early-stage and metastatic triple-negative breast cancer: a review. Hematol Oncol Clin North Am 27(4): 737–749, viii

American Cancer Society (2016) Colorectal Cancer Facts & Figures 2014-2016. Colorectal Cancer Facts & Figures publications

BoulayJL, MildG, LowyA, ReuterJ, LagrangeM, TerraccianoL, LafferU, HerrmannR, RochlitzC (2003) SMAD7 is a prognostic marker in patients with colorectal cancer. Int J Cancer 104(4): 446–449

BrierleyJD, GospondarowiczMK, WittekindC (2009) TNM Classification of Malignant Tumours, 7th Edition.

BroderickP, Carvajal-CarmonaL, PittmanAM, WebbE, HowarthK, RowanA, LubbeS, SpainS, SullivanK, FieldingS, JaegerE, VijayakrishnanJ, KempZ, GormanM, ChandlerI, PapaemmanuilE, PenegarS, WoodW, SellickG, QureshiM, TeixeiraA, DomingoE, BarclayE, MartinL, SieberO, ConsortiumC, KerrD, GrayR, PetoJ, CazierJB, TomlinsonI, HoulstonRS (2007) A genome-wide association study shows that common alleles of SMAD7 influence colorectal cancer risk. Nature genetics 39(11): 1315–1317

CunninghamD, HumbletY, SienaS, KhayatD, BleibergH, SantoroA, BetsD, MueserM, HarstrickA, VerslypeC, ChauI, Van CutsemE (2004) Cetuximab monotherapy and cetuximab plus irinotecan in irinotecan-refractory metastatic colorectal cancer. N Engl J Med 351(4): 337–345

CurryEL, MoadM, RobsonCN, HeerR (2015) Using induced pluripotent stem cells as a tool for modelling carcinogenesis. World J Stem Cells 7(2):461–469

CurtinK, LinWY, GeorgeR, KatoryM, ShortoJ, Cannon-AlbrightLA, BishopDT, CoxA, CampNJ (2009) Meta association of colorectal cancer confirms risk alleles at 8q24 and 18q21. Cancer epidemiology, biomarkers & prevention: a publication of the American Association for Cancer Research, cosponsored by the American Society of Preventive Oncology 18(2): 616–621

EpperlyMW, CaoS, ZhangX, FranicolaD, ShenH, GreenbergerEE, EpperlyLD, GreenbergerJS (2007) Increased longevity of hematopoiesis in continuous bone marrow cultures derived from NOS1 (nNOS, mtNOS) homozygous recombinant negative mice correlates with radioresistance of hematopoietic and marrow stromal cells. Exp Hematol 35(1): 137–145

FerlayJ, SoerjomataramI, DikshitR, EserS, MathersC, RebeloM, ParkinDM, FormanD, BrayF (2015) Cancer incidence and mortality worldwide: sources, methods and major patterns in GLOBOCAN 2012. Int J Cancer 136(5): E359–386

GaedckeJ, GradeM, CampsJ, SokildeR, KaczkowskiB, SchetterAJ, DifilippantonioMJ, HarrisCC, GhadimiBM, MollerS, BeissbarthT, RiedT, LitmanT (2012) The rectal cancer microRNAome-microRNA expression in rectal cancer and matched normal mucosa. Clin Cancer Res 18(18): 4919–4930

GradeM, HummonAB, CampsJ, EmonsG, SpitznerM, GaedckeJ, HoermannP, EbnerR, BeckerH, DifilippantonioMJ, GhadimiBM, BeissbarthT, CaplenNJ, RiedT (2011) A genomic strategy for the functional validation of colorectal cancer genes identifies potential therapeutic targets. Int J Cancer128(5): 1069–1079

HaimanCA, Le MarchandL, YamamatoJ, StramDO, ShengX, KolonelLN, WuAH, ReichD, HendersonBE (2007) A common genetic risk factor for colorectal and prostate cancer. Nature genetics 39(8): 954–956

HalderSK, BeauchampRD, DattaPK(2005) Smad7 induces tumorigenicity by blocking TGF-beta-induced growth inhibition and apoptosis. Exp Cell Res 307(1): 231–246

HalderSK, RachakondaG, DeaneNG, DattaPK (2008) Smad7 induces hepatic metastasis in colorectal cancer. British journal of cancer 99(6): 957–965.

HongTS, ClarkJW, HaigisKM (2012) Cancers of the colon and rectum: identical or fraternal twins? Cancer Discov 2(2):117–121

HoulstonRS, CheadleJ, DobbinsSE, TenesaA, JonesAM, HowarthK, SpainSL, BroderickP, DomingoE, FarringtonS, PrendergastJG, PittmanAM, TheodoratouE, SmithCG, OlverB, WaltherA, BarnetsonRA, ChurchmanM, JaegerEE, PenegarS, BarclayE, MartinL, GormanM, MagerR, JohnstoneE, MidgleyR, NiittymakiI, TuupanenS, ColleyJ, IdziaszczykS, ThomasHJ, LucassenAM, EvansDG, MaherER, MaughanT, DimasA, DermitzakisE, CazierJB, AaltonenLA, PharoahP, KerrDJ, Carvajal-CarmonaLG, CampbellH, DunlopMG, TomlinsonIP (2010) Meta-analysis of three genome-wide association studies identifies susceptibility loci for colorectal cancer at 1q41, 3q26.2, 12q13.13 and 20q13.33. Nature genetics 42(11): 973–977

HoulstonRS, WebbE, BroderickP, PittmanAM, BernardoMC Di, LubbeS, ChandlerI, VijayakrishnanJ, SullivanK, PenegarS, Carvajal-CarmonaL, HowarthK, JaegerE, SpainSL, WaltherA, BarclayE, MartinL, GormanM, DomingoE, TeixeiraAS, KerrD, CazierJB, NiittymakiI, TuupanenS, KarhuA, AaltonenLA, TomlinsonIP, FarringtonSM, TenesaA, PrendergastJG, BarnetsonRA, CetnarskyjR, PorteousME, PharoahPD, KoesslerT, HampeJ, BuchS, SchafmayerC, TepelJ, SchreiberS, VolzkeH, Chang-ClaudeJ, HoffmeisterM, BrennerH, ZankeBW, MontpetitA, HudsonTJ, GallingerS, CampbellH, DunlopMG (2008) Meta-analysis of genome-wide association data identifies four new susceptibility loci for colorectal cancer. Nature genetics 40(12): 1426–1435

IgnatenkoNA, HolubecH, BesselsenDG, Blohm-MangoneKA, Padilla-TorresJL, NagleRB, de AlborancIM, GuillenRJ, GernerEW (2006) Role of c-Myc in intestinal tumorigenesis of the ApcMin/+ mouse. Cancer Biol Ther 5(12): 1658–1664

JonesAM, HowarthKM, MartinL, GormanM, MihaiR, MossL, AutonA, LemonC, MehannaH, MohanH, ClarkeSE, WadsleyJ, MaciasE, CoatesworthA, BeasleyM, RoquesT, MartinC, RyanP, GerrardG, PowerD, BremmerC, ConsortiumT, TomlinsonI, Carvajal-CarmonaLG (2012) Thyroid cancer susceptibility polymorphisms: confirmation of loci on chromosomes 9q22 and 14q13, validation of a recessive 8q24 locus and failure to replicate a locus on 5q24. J Med Genet 49(3):158–163

KimT, CuiR, JeonYJ, LeeJH, LeeJH, SimH, ParkJK, FaddaP, TiliE, NakanishiH, HuhMI, KimSH, ChoJH, SungBH, PengY, LeeTJ, LuoZ, SunHL, WeiH, AlderH, OhJS, ShimKS, KoSB, CroceCM (2014) Long-range interaction and correlation between MYC enhancer and oncogenic long noncoding RNA CARLo-5. Proc Natl Acad Sci USA 111(11): 4173–8

KrauseM, DubrovskaA, LingeA, BaumannM (2017) Cancer stem cells: Radioresistance, prediction of radiotherapy outcome and specific targets for combined treatments. Adv Drug Deliv Rev 109: 63–73

LiL, PlummerSJ, ThompsonCL, MerkulovaA, AchesonLS, TuckerTC, CaseyG (2008) A common 8q24 variant and the risk of colon cancer: a population-based case-control study. Cancer epidemiology, biomarkers & prevention: a publication of the American Association for Cancer Research, cosponsored by the American Society of Preventive Oncology 17(2): 339–342

LingH, SpizzoR, AtlasiY, NicolosoM, ShimizuM, RedisRS, NishidaN, GafaR, SongJ, GuoZ, IvanC, BarbarottoE, De VriesI, ZhangX, FerracinM, ChurchmanM, van GalenJF, BeverlooBH, ShariatiM, HaderkF, EstecioMR, Garcia-ManeroG, PatijnGA, GotleyDC, BhardwajV, ShureiqiI, SenS, MultaniAS, WelshJ, YamamotoK, TaniguchiI, SongMA, GallingerS, CaseyG, ThibodeauSN, Le MarchandL, TiirikainenM, ManiSA, ZhangW, DavuluriRV, MimoriK, MoriM, SieuwertsAM, MartensJW, TomlinsonI, NegriniM, Berindan-NeagoeI, FoekensJA, HamiltonSR, LanzaG, KopetzS, FoddeR, CalinGA (2013) CCAT2, a novel noncoding RNA mapping to 8q24, underlies metastatic progression and chromosomal instability in colon cancer. Genome Res 23(9): 1446–1461

LuoL, LiN, LvN, HuangD (2014) SMAD7: a timer of tumor progression targeting TGF-beta signaling. Tumour Biol 35(9):8379–8385

PetersU, BienS, ZubairN(2015) Genetic architecture of colorectal cancer. Gut 64(10): 1623–1636

PetersU, JiaoS, SchumacherFR, HutterCM, AragakiAK, BaronJA, BerndtSI, BezieauS, BrennerH, ButterbachK, CaanBJ, CampbellPT, CarlsonCS, CaseyG, ChanAT, Chang-ClaudeJ, ChanockSJ, ChenLS, CoetzeeGA, CoetzeeSG, ContiDV, CurtisKR, DugganD, EdwardsT, FuchsCS, GallingerS, GiovannucciEL, GogartenSM, GruberSB, HaileRW, HarrisonTA, HayesRB, HendersonBE, HoffmeisterM, HopperJL, HudsonTJ, HunterDJ, JacksonRD, JeeSH, JenkinsMA, JiaWH, KolonelLN, KooperbergC, KuryS, LacroixAZ, LaurieCC, LaurieCA, Le MarchandL, LemireM, LevineD, LindorNM, LiuY, MaJ, MakarKW, MatsuoK, NewcombPA, PotterJD, PrenticeRL, QuC, RohanT, RosseSA, SchoenRE, SeminaraD, ShrubsoleM, ShuXO, SlatteryML, TavernaD, ThibodeauSN, UlrichCM, WhiteE, XiangY, ZankeBW, ZengYX, ZhangB, ZhengW, HsuL (2013) Identification of Genetic Susceptibility Loci for Colorectal Tumors in a Genome-Wide Meta-analysis. Gastroenterology 144(4): 799–807.e24

PomerantzMM, AhmadiyehN, JiaL, HermanP, VerziMP, DoddapaneniH, BeckwithCA, ChanJA, HillsA, DavisM, YaoK, KehoeSM, LenzHJ, HaimanCA, YanC, HendersonBE, FrenkelB, BarretinaJ, BassA, TaberneroJ, BaselgaJ, ReganMM, ManakJR, ShivdasaniR, CoetzeeGA, FreedmanML (2009) The 8q24 cancer risk variant rs6983267 shows long-range interaction with MYC in colorectal cancer. Nature genetics 41(8): 882–884

RajagopalanMS, StoneB, RwigemaJC, SalimiU, EpperlyMW, GoffJ, FranicolaD, DixonT, CaoS, ZhangX, BuchholzBM, BauerAJ, ChoiS, BakkenistC, WangH, GreenbergerJS (2010) Intraesophageal manganese superoxide dismutase-plasmid liposomes ameliorates novel total-body and thoracic radiation sensitivity of NOS1-/- mice. Radiat Res 174(3): 297–312

RennollS, YochumG (2015) Regulation of MYC gene expression by aberrant Wnt/beta-catenin signaling in colorectal cancer. World J Biol Chem 6(4): 290–300

RiihimakiM, HemminkiA, SundquistJ, HemminkiK (2016) Patterns of metastasis in colon and rectal cancer. Sci Rep 6: 29765

RodelC, HofheinzR, LierschT (2012a) Rectal cancer: state of the art in 2012. Curr Opin Oncol 24(4):441–447

RodelC, LierschT, BeckerH, FietkauR, HohenbergerW, HothornT, GraevenU, ArnoldD, Lang-WelzenbachM, RaabHR, SulbergH, WittekindC, PotapovS, StaibL, HessC, Weigang-KohlerK, GrabenbauerGG, HoffmannsH, LindemannF, Schlenska-LangeA, FolprechtG, SauerR, German Rectal Cancer Study G (2012b) Preoperative chemoradiotherapy and postoperative chemotherapy with fluorouracil and oxaliplatin versus fluorouracil alone in locally advanced rectal cancer: initial results of the German CAO/ARO/AIO-04 randomised phase 3 trial. Lancet Oncol 13(7): 679–687

RosseSA, AuerPL, CarlsonCS (2014) Functional annotation of putative regulatory elements at cancer susceptibility Loci. Cancer Inform 13(Suppl 2):5–17

SansomOJ, MenielVS, MuncanV, PhesseTJ, WilkinsJA, ReedKR, VassJK, AthineosD, CleversH, ClarkeAR (2007) Myc deletion rescues Apc deficiency in the small intestine. Nature 446(7136): 676–679

SauerR, BeckerH, HohenbergerW, RodelC, WittekindC, FietkauR, MartusP, TschmelitschJ, HagerE, HessCF, KarstensJH, LierschT, SchmidbergerH, RaabR, German Rectal Cancer StudyG (2004) Preoperative versus postoperative chemoradiotherapy for rectal cancer. N Engl J Med 351(17): 1731–1740

SmithJJ, DeaneNG, WuF, MerchantNB, ZhangB, JiangA, LuP, JohnsonJC, SchmidtC, BaileyCE, EschrichS, KisC, LevyS, WashingtonMK, HeslinMJ, CoffeyRJ, YeatmanTJ, ShyrY, BeauchampRD (2010) Experimentally derived metastasis gene expression profile predicts recurrence and death in patients with colon cancer. Gastroenterology 138(3):958–968

SpitznerM, RoeslerB, BielfeldC, EmonsG, GaedckeJ, WolffHA, Rave-FrankM, KramerF, BeissbarthT, KitzJ, WienandsJ, GhadimiBM, EbnerR, RiedT, GradeM (2014) STAT3 inhibition sensitizes colorectal cancer to chemoradiotherapy in vitro and in vivo. Int J Cancer 134(4):997–1007

TenesaA, FarringtonSM, PrendergastJG, PorteousME, WalkerM, HaqN, BarnetsonRA, TheodoratouE, CetnarskyjR, CartwrightN, SempleC, ClarkAJ, ReidFJ, SmithLA, KavoussanakisK, KoesslerT, PharoahPD, BuchS, SchafmayerC, TepelJ, SchreiberS, VolzkeH, SchmidtCO, HampeJ, Chang-ClaudeJ, HoffmeisterM, BrennerH, WilkeningS, CanzianF, CapellaG, MorenoV, DearyIJ, StarrJM, TomlinsonIP, KempZ, HowarthK, Carvajal-CarmonaL, WebbE, BroderickP, VijayakrishnanJ, HoulstonRS, RennertG, BallingerD, RozekL, GruberSB, MatsudaK, KidokoroT, NakamuraY, ZankeBW, GreenwoodCM, RangrejJ, KustraR, MontpetitA, HudsonTJ, GallingerS, CampbellH, DunlopMG (2008) Genome-wide association scan identifies a colorectal cancer susceptibility locus on 11q23 and replicates risk loci at 8q24 and 18q21. Nature genetics 40(5): 631–637

ThompsonCL, PlummerSJ, AchesonLS, TuckerTC, CaseyG, LiL (2009) Association of common genetic variants in SMAD7 and risk of colon cancer. Carcinogenesis30(6):982–986

TomlinsonI, WebbE, Carvajal-CarmonaL, BroderickP, KempZ, SpainS, PenegarS, ChandlerI, GormanM, WoodW, BarclayE, LubbeS, MartinL, SellickG, JaegerE, HubnerR, WildR, RowanA, FieldingS, HowarthK, ConsortiumC, SilverA, AtkinW, MuirK, LoganR, KerrD, JohnstoneE, SieberO, GrayR, ThomasH, PetoJ, CazierJB, HoulstonR (2007) A genome-wide association scan of tag SNPs identifies a susceptibility variant for colorectal cancer at 8q24.21. Nature genetics 39(8): 984–988

TuupanenS, TurunenM, LehtonenR, HallikasO, VanharantaS, KiviojaT, BjorklundM, WeiG, YanJ, NiittymakiI, MecklinJP, JarvinenH, RistimakiA, Di-BernardoM, EastP, Carvajal-CarmonaL, HoulstonRS, TomlinsonI, PalinK, UkkonenE, KarhuA, TaipaleJ, AaltonenLA (2009) The common colorectal cancer predisposition SNP rs6983267 at chromosome 8q24 confers potential to enhanced Wnt signaling. Nature genetics 41(8): 885–890

van der SijpMP, BastiaannetE, MeskerWE, van der GeestLG, BreugomAJ, SteupWH, MarinelliAW, TsengLN, TollenaarRA, van de VeldeCJ, DekkerJW (2016) Differences between colon and rectal cancer in complications, short-term survival and recurrences. Int J Colorectal Dis 31(10): 1683–1691

VerziMP, HatzisP, SulahianR, PhilipsJ, SchuijersJ, ShinH, FreedE, LynchJP, DangDT, BrownM, CleversH, LiuXS, ShivdasaniRA (2010) TCF4 and CDX2, major transcription factors for intestinal function, converge on the same cis-regulatory regions. Proc Natl Acad Sci U S A 107(34): 15157–15162

WhiffinN, HoskingFJ, FarringtonSM, PallesC, DobbinsSE, ZgagaL, LloydA, KinnersleyB, GormanM, TenesaA, BroderickP, WangY, BarclayE, HaywardC, MartinL, BuchananDD, WinAK, HopperJ, JenkinsM, LindorNM, NewcombPA, GallingerS, ContiD, SchumacherF, CaseyG, LiuT, CampbellH, LindblomA, HoulstonRS, TomlinsonIP, DunlopMG (2014) Identification of susceptibility loci for colorectal cancer in a genome-wide metaanalysis. Human molecular genetics 23(17): 4729–4737

WrightJB, BrownSJ, ColeMD (2010) Upregulation of c-MYC in cis through a large chromatin loop linked to a cancer risk-associated single-nucleotide polymorphism in colorectal cancer cells. Mol Cell Biol 30(6): 1411–1420

YeagerM, OrrN, HayesRB, JacobsKB, KraftP, WacholderS, MinichielloMJ, FearnheadP, YuK, ChatterjeeN, WangZ, WelchR, StaatsBJ, CalleEE, FeigelsonHS, ThunMJ, RodriguezC, AlbanesD, VirtamoJ, WeinsteinS, SchumacherFR, GiovannucciE, WillettWC, Cancel-TassinG, CussenotO, ValeriA, AndrioleGL, GelmannEP, TuckerM, GerhardDS, FraumeniJFJr., HooverR, HunterDJ, ChanockSJ, ThomasG (2007) Genome-wide association study of prostate cancer identifies a second risk locus at 8q24. Nature genetics 39(5): 645–649

ZhangK, CivanJ, MukherjeeS, PatelF, YangH (2014) Genetic variations in colorectal cancer risk and clinical outcome. World journal of gastroenterology: WJG 20(15): 4167–4177

ZhuRX, SongCH, YangJS, YiQT, LiBJ, LiuSH (2016) Downregulation of AATK mediates microRNA-558- induced resistance of A549 cells to radiotherapy. Mol Med Rep14(3):2846–2852

